# Selective BCL-X_L_ Antagonists Eliminate Infected Cells from a Primary Cell Model of HIV Latency but not from *Ex Vivo* Reservoirs

**DOI:** 10.1101/2020.11.25.397679

**Authors:** Yanqin Ren, Szu Han Huang, Amanda B. Macedo, Adam R. Ward, Winiffer D. Conce Alberto, Thais Klevorn, Louise Leyre, Dora Chan, Ronald Truong, Thomas Rohwetter, Paul Zumbo, Friederike Dündar, Doron Betel, Colin Kovacs, Erika Benko, Alberto Bosque, R. Brad Jones

## Abstract

HIV persists, despite antiviral immune responses and effective antiretroviral therapy, in viral reservoirs that seed rebound viremia if therapy is interrupted. Previously, we showed that the BCL-2 protein contributes to HIV persistence by conferring a survival advantage to reservoir-harboring cells. Here, we demonstrate that many of the BCL-2 family members are overexpressed in HIV-infected CD4^+^ T-cells, indicating increased tension between pro-apoptotic and pro-survival family members – as well as raising the possibility that the inhibition of pro-survival members may disproportionately affect the survival of HIV-infected cells. Based on these results, we chose to further study *BCL2L1* (encoding the protein BCL-X_L_), due to its consistent overexpression and the availability of selective antagonists. Infection of primary CD4^+^ T-cells with either a clinical isolate, a CCR5-tropic strain, or a CXCR4-tropic strain of HIV resulted in increased BCL-X_L_ protein expression; and treatment with two selective BCL-X_L_ antagonists, A-1155463 and A-1551852, led to disproportionate cell death compared to uninfected CD4^+^ T-cells. In a primary cell model of latency, both BCL-X_L_ antagonists drove significant reductions in total HIV DNA and in infectious cell frequencies both alone and in combination with the latency reversing agent bryostatin-1, with little off-target cytotoxicity. However, these antagonists, with or without bryostatin-1, or in combination with the highly potent latency reversing agent combination PMA + ionomycin, failed to reduce total HIV DNA and infectious reservoirs in *ex vivo* CD4^+^ T-cells from ART-suppressed donors. Our results add to growing evidence that bonafide reservoir-harboring cells are resistant to multiple “kick and kill” modalities - relative to latency models - and uncover BCL-X_L_ antagonists as a facile approach to probing mechanistic underpinnings. We also interpret our results as encouraging of further exploration of BCL-X_L_ antagonists for cure, where combination approaches may unlock the ability to eliminate *ex vivo* reservoirs.

## Introduction

Although antiretroviral therapy (ART) durably suppresses HIV replication, it cannot eradicate persisting infected cells with integrated HIV proviruses. These cells comprise a ‘viral reservoir’ which seeds viral rebound when ART is interrupted, with the best characterized reservoir being formed in CD4^+^ T-cells [1]. An important mechanism of persistence is the maintenance of viral latency, particularly in resting memory CD4^+^ T-cells, which prevents death from viral cytopathic effects as well as recognition and elimination by immune effectors, such as cytotoxic T-lymphocytes (CTLs) [2–4]. More recent work has additionally uncovered mechanisms by which reservoir-harboring cells may survive both viral- or immune-mediated cytopathicity, with the pro-survival factor BCL-2 implicated in both of these scenarios [5]. With respect to viral cytopathicity, one mechanism of death is through cleavage of the host protein procaspase 8 by HIV protease to generate Casp8p41, which drives apoptosis [6]. BCL-2 is able to antagonize this pathway by binding to Casp8p41, and preventing cell death in BCL-2^high^ cells [7]. With respect to immune-mediated cytopathicity, our group recently made a series of observations which led us to conclude that BCL-2 comprises one mechanism by which HIV reservoir-harboring cells resist elimination by CTL: i) BCL-2^high^ CD4^+^ T-cells preferentially survive CTL killing *in vitro*, ii) the inducible HIV reservoir in *ex vivo* CD4^+^ T-cells is disproportionately harbored in BCL-2^high^ CD4^+^ T-cells, and iii) the BCL-2 antagonist ABT-199 sensitizes *ex vivo* HIV reservoirs to reductions by combinations of HIV-specific T-cells and latency reversing agents (LRAs) [8]. While much study to date has focused on BCL-2, it is likely that this is only one of the mechanisms that govern the intrinsic susceptibility of an HIV reservoir-harboring cell to death. This idea draws support from a recent study which demonstrated a role for BIRC5, also known as survivin, in activating cellular survival programs to promote HIV persistence - BIRC5 was found to be over-expressed in HIV reservoir-harboring cells that had undergone clonal expansion [9]. Inhibition of this protein resulted in a selective decrease of HIV-infected cells, suggesting that BIRC5 supports long-term survival of HIV-infected cells [9]. Building upon these results, the fundamental premise of the current study is that additional pro-survival mechanisms remain to be discovered, with our current focus being on those which protect cells from viral cytopathic effects. We focus here on BCL-2 family members, as a logical extenson of the current state of knowledge.

BCL-2 is the prototypical member of a family of proteins which define the pro- or anti-apoptotic states of a cell through their complex interplay (reviewed in [10]). The pro-death members (e.g. BAX) are activated in response to a range of deleterious events, and act to form channels which allow cytochrome c to exit the mitochondria, which activates caspases and induces cell death [11, 12]. The pro-survival proteins (e.g. BCL-2) inhibit those pro-apoptotic partners, leading to the balance of the original rheostat model [13–15]. The expression of a number of BCL-2 family members have been reported to be modulated by HIV infection and replication, though their potential roles in HIV persistence remain largely unstudied [16]. The expression of the pro-survival member BCL-2 has been reported to be reduced in CD4^+^ T-cells by gp120 cross-linking, which induces cell apoptosis [17]. HIV-Env can also elicit p53-dependent Puma, followed by activating Bax and Bak to induce apoptosis [18]. The HIV Tat protein has been implicated in the upregulation of a number of pro-apoptotic members, including the proteins Bim, Puma, and Noxa, through either activation of the FOXO3a transcriptional activator, or downregulation of BCL-2 expression, resulting in apoptosis through mitochondrial membrane permeabilization [19–21]. HIV-Vpr can bind to adenine nucleotide translocase (ANT), a protein that forms the inner membrane channel of the mitochondrial permeability transition pore (MPTP), then converts it into a pro-apoptotic pore, which leads to cell death [22, 23]. Also acting to promote apoptosis, the pro-survival BCL-2 homologs BCL-X_L_ and Bfl1/A1 have been reported to be suppressed by HIV Vpu [24]. On the pro-survival side of the interaction, the pro-apoptotic homolog BAD has been shown to be inactivated by HIV Nef, through phosphorylation [25]. Low level HIV-Vpr expression on Jurkat cells can upregulate BCL-2 and downmodulate BAX, facilitating HIV persistence [26]. HIV-Tat and gp120 can induce TREM-1 expression in macrophages, and TREM-1 leads to inactivation of caspase 3 and increased BCL-2 expression, thus inhibiting apoptosis [27]. On the systemic level, in PBMCs from people living with HIV, BCL-2, BCL-X_L_ and MCL‐1 were significantly upregulated during successful ART [28]. Puma expression was enhanced in untreated HIV-infected donors, and these elevated Puma levels decreased upon ART initiation [18]. Overall, these results paint a nuanced picture of apoptotic signaling in the setting of HIV infection, though despite these individual reports of BCL-2 family member modulation – often in the context of cell lines – the overall impact of BCL-2 family proteins on infected cell fate is not completely understood.

Here, we sought to define the landscape of BCL-2 family expression during HIV infection, with the aim of identifying a targetable factor that may be important to HIV persistence. Transcriptional profiling of HIV-infected versus uninfected cells from *in vitro* primary CD4^+^ T-cell infection cultures revealed that multiple BCL-2 family members were significantly differentially expressed, including both pro-survival and pro-apoptotic actors. We selected BCL-X_L_ (*BCL2L1*), a pro-survival factor [29], for futher study based on its consistent upregulation in HIV-infected cells, and the availability of selective inhibitors under drug development for cancer. Overexpresson of BCL-X_L_ in HIV-infected cells was confirmed at the protein level by flow cytometry, and we observed that two selective BCL-X_L_ inhibitors enhanced the death of productively HIV-infected cells *in vitro*. Both inhibitors also consistently drove the selective death of HIV-infected cells from primary cell models of latency, either with or without the LRA bryostatin – with very little bystander toxicity. However, when we applied the same strategies to ‘natural’ HIV reservoirs in *ex vivo* CD4^+^ T-cells from ARV-treated donors, we observed that none of these treatments were sufficient to reduce reservoir sizes. Our study thus identifies BCL-X_L_ overexpression as a survival mechanism that can be targeted in productive-infection to promote death of the HIV-infected cells, and also possibly in a latency-reactivation setting, though it remains to be determined if combining BCL-X_L_ antagonism with immune effectors or other modes of infected cell killing can achieve a reduction in *ex vivo* reservoirs. The discordant results beween a primary-cell latency model versus *ex vivo* HIV reservoirs (in closely matched experimental conditions) parallels our previous observations with CTL-based ‘kick and kill’ combinations [8], or to a fundamental resistance of *ex vivo* reservoirs to multiple modes of attack. Our study identifies BCL-X_L_ antagonists as a novel and facile ‘kill’ component (without the bystander toxicity of BCL-2 antagonists) that can be leveraged to study the survivability of *ex vivo* reservoirs. The identification of additional barriers to the elimination of these reservoirs may yield combination treatment strategies which enable the selective toxicity of BCL-X_L_ antagonists to HIV-infected cells observed in productive infection and latency models to be brought to bear against reservoir-harboring cells.

## Results

### Transcriptional profiling of BCL-2 family members reveals increased tension in HIV-infected vs uninfected cells

To test whether expression of BCL-2 family proteins are altered during HIV infection, we infected CD4^+^ T-cells from a long-term ARV-suppressed donor ‘OM5267’ with the HIV virus ‘JR-CSF’, and assessed the expression of BCL-2 family transcripts by RNA sequencing (RNAseq). In an initial experiment, we used total CD4^+^ T-cells for infection, and sorted them into HIV-positive (Gag^+^) and HIV-negative (Gag^−^) using flow cytometry (**Fig. 1A)**. Of all the genes that we compared, we observed a cluster of BCL-2 family member s[30], (22 genes) that were significantly upregulated in HIV-positive cells, which included the pro-apoptotic members *BAK1* (encodes the protein BAK), *BMF*, *PMAIP1* (encodes the protein NOXA) and *BBC3* (encodes the protein PUMA), and the pro-survival members *BCL2L1* (encodes BCL-X_L_) and *BCL2A1* (encodes BCL-2-related protein A1) (**Fig. 1B**). Only two BCL-2 family genes were significantly under-expressed in HIV-positive cells: the pro-survival member *MCL1* and pro-apoptotic member *BAX*. Thus, HIV-infected primary CD4^+^ T-cells were predominately characterized by overexpression of the BCL-2 family, representing both pro-survival and pro-apoptotic members.

**Fig. 1.**
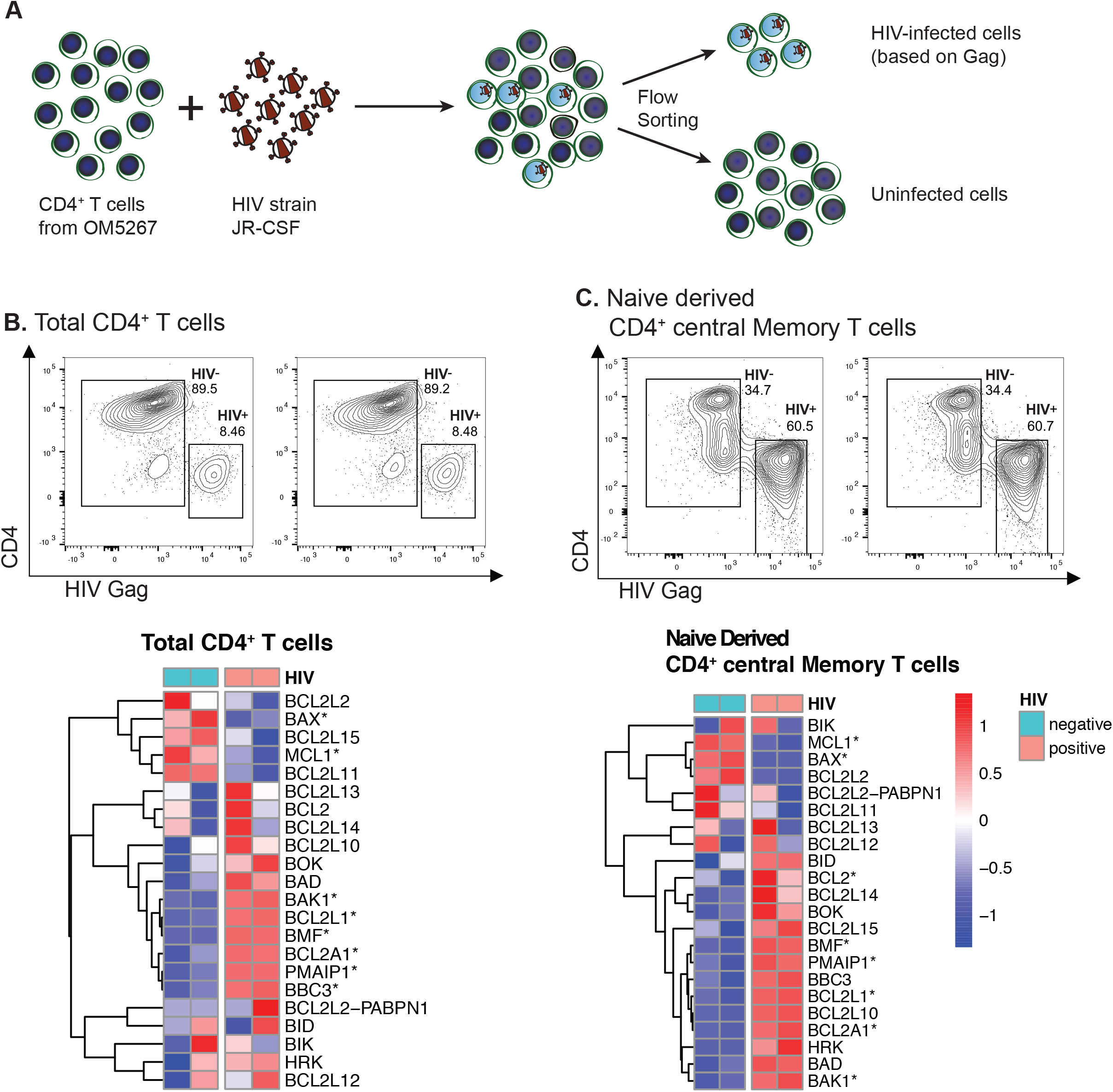
Transcriptional profiles of BCL-2 family proteins reveal differences in expression of pro-survival and pro-apoptotic genes between HIV-infected cells and uninfected cells. **(A)** Schematic of HIV infection (*in vitro*) and flow cytometry sorting based on intracellular HIV-p24 (Gag) expression: Gag^+^ represents HIV-positive cells, Gag^−^ represents HIV-negative cells. **(B-C)** Flow gating strategies for cell sorting and corresponding expression patterns of BCL-2 family genes in HIV-positive vs negative cells. Samples were derived from total CD4^+^ T cells (**B**) or from Naïve CD4^+^ T cells differentiated into central memory T cells (**C**) that were then infected with HIV *in vitro*. The heatmaps visualize the scaled expression values based on RNA-seq of two technical replicates of HIV-positive and HIV-negative cells (separate infection and sorting). Asterisks (*) highlight the genes that were found to be significantly differentially expressed between HIV-infected cells vs. uninfected cells (padj<0.01).

The above experiment utilized total CD4^+^ T-cells and thus contained cells of diverse phenotypes and functional profiles. We next extended these results to an analogous experiment using a more homogeneous pool of cells with a central memory (T_CM_) phenotype, given that these cells are a major cellular reservoir of HIV [31]. T_CM_ were generated by the *in vitro* priming of naïve CD4^+^ T-cells, using the methodology of the well-characterized cultured T_CM_ model of HIV latency [32, 33]. With this more homogeneous cell population, we again observed a general pattern of overexpression of BCL-2 family genes in HIV-infected vs uninfected cells, including the pro-apoptotic members *PMAIP1*, *BMF*, and *BAK1*, and the pro-survival members *BCL2*, *BCL2A1* and *BCL2L1* (**Fig. 1C**). As with total CD4^+^ T-cells, we observed significant underexpression of the pro-survival gene *MCL1* in HIV-positive cells and the pro-apoptotic member *BAX*. Together, these data indicate an increased tension between pro-apoptotic and pro-survival BCL-2 family members in HIV-infected cells as compared to uninfected cells, suggesting to us that the inhibition of pro-survival members may disproportionately lead to the death of HIV-infected cells by tipping the balance in favor of the pro-apoptotic members that are overexpressed in these cells. While the data propose several pro-survival members that could be targeted to test this hypothesis, we opted here to focus on BCL-X_L_, in part due to the availability of selective antagonists.

### Selective BCL-X_L_ antagonists disproportionately eliminate HIV infected cells

In order to evaluate if BCL2L1 overexpression translates to increased BCL-X_L_ protein expression in HIV-infected vs uninfected cells, protein levels were measured using flow cytometry. Activated CD4^+^ T cells, isolated from an HIV-negative donor, were infected with 3 different viruses – one clinical isolate from a quantitative viral outgrowth assay (QVOA virus), one CCR5-tropic strain (JR-CSF), and one CXCR4-tropic strain (NL4-3). One week later, infected cells were treated with a BCL-X_L_ antagonist, either A-1155463 or A-1331852, or DMSO (negative control), under the suppression of T-20 (an HIV fusion inhibitor) in order to prevent new rounds of infection during the treatments. 48 hours later, cells were stained with live/dead and anti-BCL-X_L_ antibody to measure BCL-X_L_ expression levels and for the potential elimination of HIV-infected cells. Applying the flow cytometry gating strategy shown in **Fig. 2A**, we observed that the BCL-X_L_ MFI in HIV-infected cells (HIV-Gag^+^) was 1.86-2.25 fold higher than that in uninfected cells (HIV-Gag^−^) (**Fig. 2B**), suggesting that the virus-host interaction during the HIV infection upregulates BCL-X_L_ expression.

**Fig. 2.**
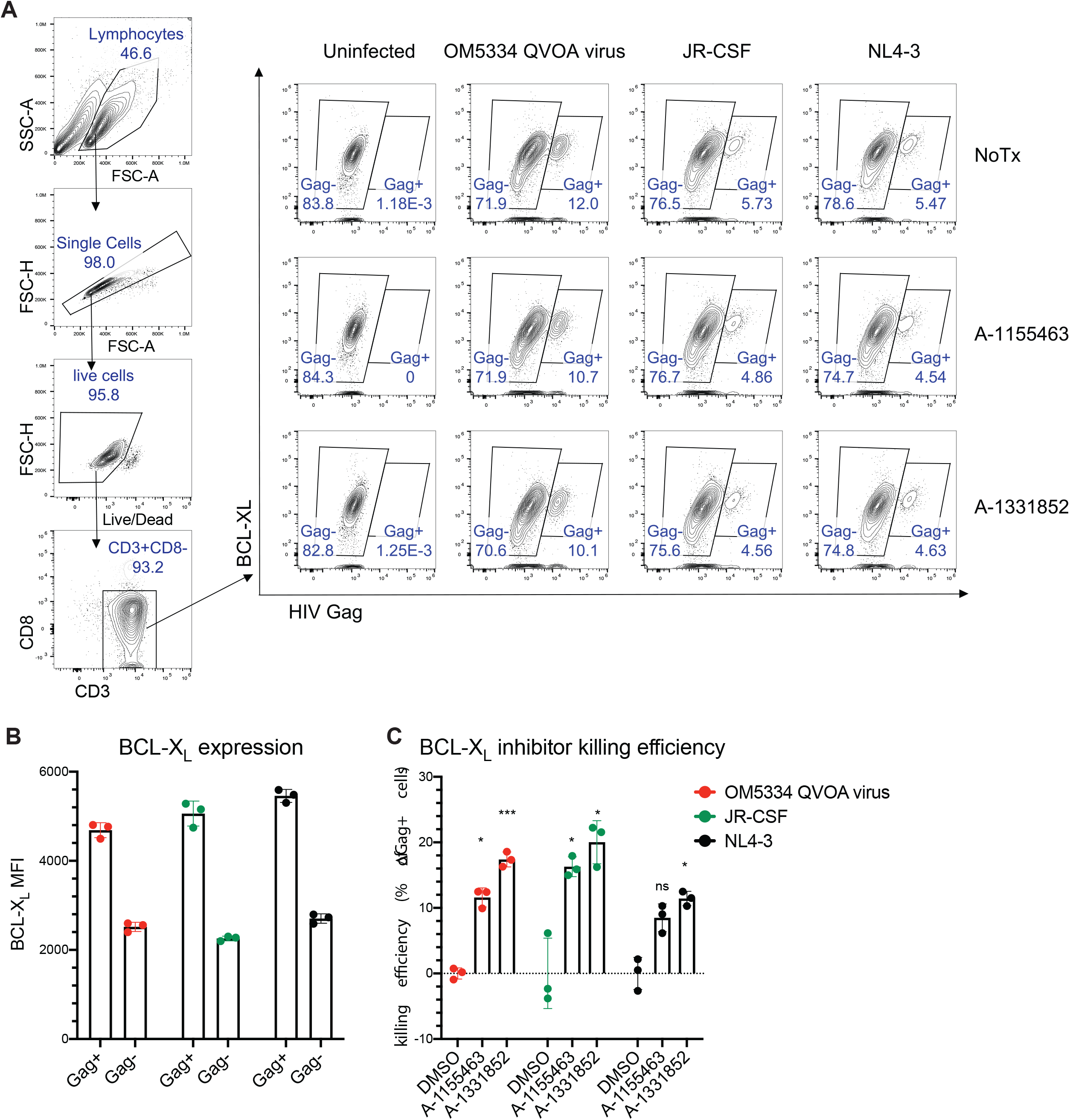
BCL-X_L_ antagonists eliminate HIV-infected cells in productive infection. **(A)** Flow cytometry gating strategy for HIV infection with 3 different viruses: clinical isolate (QVOA virus), CCR5-tropic (JR-CSF), and CXCR4-tropic (NL4-3). Two BCL-X_L_ antagonists were used for treatments (A-1155463 and A-1331852). (**B**) BCL-X_L_ expression levels (MFI) in HIV-infected cells (HIV-Gag^+^) and HIV-uninfected cells (HIV-Gag^−^). (**C**) BCL-X_L_ antagonist treatments eliminate HIV-infected cells. Killing efficiency was calculated by the following formula: (% p24^+^ in DMSO condition minus % p24^+^ in BCL-X_L_ antagonist condition)/ (% p24^+^ in DMSO condition). Statistical significance was determined by two-way ANOVA, using Dunnett's multiple comparisons test. * padj<0.05, ** padj<0.01, *** padj<0.001, **** padj<0.0001.

To test if HIV-infected cells could be preferentially eliminated by the two selective BCL-X_L_ antagonists, A-1155463 and A-1331852, we measured the killing efficiency by counting on the Gag^+^ cells percentage in each treatment and comparing this to DMSO treatment (negative control). Cell death was observed disproportionally in HIV-infected cells, where we showed a 12.12% average decrease in the percentage of HIV-infected cells relative to DMSO control for A-1155463 (11.60%, 16.26% and 8.49% for OM5334 QVOA virus, JR-CSF and NL4-3, respectively), and a 16.27% average decrease for A-1331852 (17.40%, 20.00% and 11.42% for OM5334 QVOA virus, JR-CSF and NL4-3, respectively) (**Fig. 2C**). These results suggest that BCL-X_L_ is overexpressed in HIV-infected cells, and that the selective BCL-X_L_ antagonists are able to selectively eliminate some propotion of productively HIV-infected cells.

### Selective BCL-X_L_ antagonists drive reductions in total HIV DNA and infectious reservoirs in a primary cell latency model

We next used the two selective BCL-X_L_ antagonists to test the hypothesis that BCL-X_L_ antagonism could sensitize HIV reservoir-harboring cells to elimination. This mirrors previous work in which we reported that the BCL-2 antagonist ABT-199 sensitized HIV reservoir-harboring cells to elimination in a primary cell latency model, when used alone or in combination with a latency reversing agent [supplementary data of [8]). The primary cell model of HIV latency (see methods for cell generation details) typically gives rise to frequencies of latently-infected cells between ~2%-5%, greatly exceeding the ~0.0001% that is typical of *ex vivo* CD4^+^ T-cells from ARV-treated individuals which harbor intact-inducible virus. The QVOA is the gold standard method for quantifying the inducible reservoir, and it is well-suited for rare *ex vivo* populations [34], however it cannot be directly applied to the much higher frequencies of infected cells in the primary cell latency model. To set the stage for direct comparisons between results from the primary cell latency model and those of *ex vivo* reservoirs, we therefore spiked these model cells into autologous CD4^+^ T-cells in order to bring the frequency of latently infected cells into closer alignment with *ex vivo* reservoirs (aiming for ~1,000–10,000 copies of HIV DNA per million CD4^+^ T-cells). The effects of the selective BCL-X_L_ antagonists were then assessed using an HIV eradication (HIVE) assay (**Fig. 3A**). HIVE assays consist of a 4-day culture with one of the selective BCL-X_L_ antagonists in the presence of ARVs, with or without prior activation of target cells with the latency reversing agent bryostatin-1 (a protein kinase C agonist), which is washed out prior to the culture period. Surviving cells are then counted and subjected to droplet digital PCR (ddPCR) to measure total frequencies of infected cells, and QVOA to measure intact-inducible reservoirs. This distinction is important as, in *ex vivo* CD4^+^ T-cells from ARV-treated individuals, the large majority of HIV DNA represents defective proviruses with no potential for viral replication [34]. In contrast to the BCL-2 antagonist ABT-199, which was associated with substantial levels of cell death, the selective BCL-X_L_ antagonists A-1155463 and A-1331852 showed little or no cytotoxicity at 100nM (**SFig. 1**). We were nonetheless careful to account for cell death in our QVOA assays by counting only viable CD4^+^ T-cells by flow cytometry following a 24-hour drug wash-out period, and using only viable cell numbers to calculate infectious units per million CD4^+^ T-cells (IUPM).

**Fig. 3.**
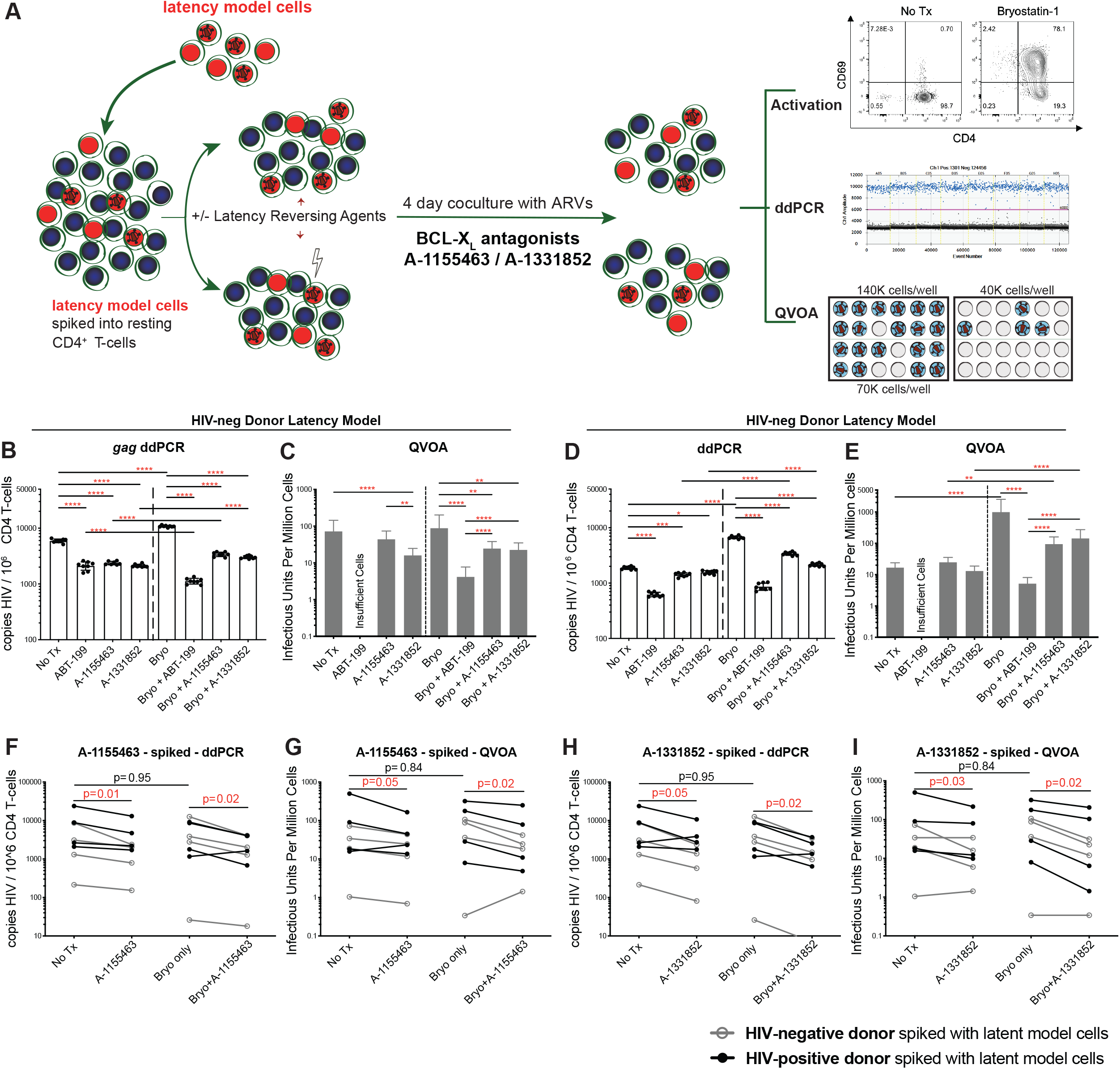
BCL-X_L_ antagonists drive reductions in total HIV DNA and infectious reservoirs in a primary cell model of latency in “spiked” HIVE assays. **(A)** Schematic of a HIVE assay “spiked" with a primary cell latency model. **(B-C)** Representative “spiked” HIVE assay with cells from an <u>HIV-uninfected donor</u>. **(B)** ddPCR results showing the mean HIV DNA levels ± SD of 8 technical replicates (HIV DNA measured with the 5’ end primer/probe set targeting HIV *gag*), **(C)** QVOA results showing IUPM ± 95% confidence interval. **(D-E)** Representative spiked HIVE assay with cells from an <u>HIV-infected donor</u>. (**D)** ddPCR results showing the mean ± SD of 8 technical replicates. **(E)** QVOA results showing IUPM ± 95% confidence interval. Statistical significance determined by: one-way ANOVA for ddPCR, and pairwise Chi-square tests for QVOA. * p<0.05, ** p<0.01, *** p<0.001, **** p<0.0001. **(F-I) Summary data for A-1155463 and A-1331852. (F)** Summary ddPCR results showing mean (of 8 replicates each) of levels of HIV DNA and **(G)** summary QVOA results showing intact inducible replication-competent proviruses (IUPM), comparing treatment with A-1155463 vs No Tx, and Bryostatin-1+A-1155463 vs Bryostatin-1 (n=8). **(H)** Levels of HIV DNA and **(I)** IUPM, comparing treatment with A-1331852 vs No Tx, and Bryostatin-1+A-1331852 vs Bryostatin-1 (n=8). Black lines with dots represent the primary cell model of latency generated with ART-suppressed HIV-infected donors, grey lines with open circles represent latency model generated with HIV-uninfected donors. DMSO was added to NoTx conditions at a matched concentration with +Tx conditions. Statistical significance was determined by Wilcoxon matched-pairs signed rank test.

In an initial experiment using model cells generated from an HIV-negative donor, we observed significant reductions in total HIV DNA as measured by *gag* primer/probe sets following treatment with each of the BCL-X_L_ antagonists alone, with a 2.50-fold average reduction for A-1155463 and a 2.79-fold average reduction for A-1331852 (**Fig. 3B**). Unexpectedly, latency reversal was not strictly required for these reductions. The BCL-2 antagonist ABT-199 was used as a positive control in this experiment (**Fig. 3B**). As assessed by total HIV DNA measurement, combination treatments with bryostatin-1 and BCL-X_L_ antagonists (A-1155463 or A-1331852) significantly reduced HIV DNA relative to bryostatin-1 alone (3.17-fold and 3.57-fold, respectively), however did not reduce HIV DNA relative to BCL-X_L_ antagonists alone (**Fig. 3B**). In this initial experiment, treatment with A-1331852 was observed to drive a significant reduction in IUPM when used alone (4.52-fold, p<0.0001, **Fig. 3C**). A-1155463 alone also showed a 1.65-fold reduction in IUPM, but this was not statistically significant. Significant reductions in IUPM were also observed in treatments with each of the BCL-X_L_ antagonists + bryostatin-1 (p<0.01), where A-1155463 + bryostatin-1 showed a 3.59-fold decrease and A-1331852 + bryostatin-1 showed a 3.90-fold decrease relative to bryostatin-1 alone, although these reductions were smaller in magnitude than that observed with ABT-199, which showed a 21.18-fold decrease relative to bryostain-1 alone (**Fig. 3C**). Thus, these data suggest that both the BCL-X_L_ antagonists were able to reduce the frequency of latently HIV-infected cells generated from an HIV-negative donor, either alone or in combination with bryostatin-1.

To account for potential differences in CD4^+^ T-cell function from HIV-negative vs people living with HIV (PLWH), we also performed HIVE assays with latency model cells generated from an ART-suppressed HIV-positive donor, spiked into autologous resting CD4^+^ T-cells [35]. As with cells from HIV-negative donors, significant reductions were observed in total HIV DNA following treatment with either of the BCL-X_L_ antagonists or ABT-199 relative to no treatment (p<0.001, **Fig. 3D**). The effects of the BCL-X_L_ antagonists alone were more modest than ABT-199, with a reduction in HIV DNA of 3.01-fold, 1.32-fold and 1.20-fold for ABT-199, A-1155463 and A-1331852, respectively (**Fig. 3D**). Combinations of either the BCL-X_L_ antagonists or ABT-199 with bryostatin-1 resulted in significant and consistent reductions in HIV DNA relative to bryostatin-1 only, with reductions of 7.72-fold, 1.97-fold and 3.12-fold, respectively (p<0.001, **Fig. 3D**). We did not observe significant decreases in IUPM for BCL-X_L_ antagonists treated alone or in combination with bryostatin-1 (**Fig. 3E**). However, we did observe a 21.18-fold significant reduction in IUPM in the bryostatin-1 + ABT-199 condition (p<0.0001, **Fig. 3E**). Thus, results using cells from this HIV-positive donor mirrored those from the HIV-negative donor, with both showing that the two BCL-X_L_ antagonists were effective in reducing infected cell frequencies in this spiked primary cell latency model, although the reduction in IUPM did not reach statistical significance in the model generated from the HIV-positive donor whereas it did in the model generated from the HIV-negative donor.

To assess the generalizability of our results with BCL-X_L_ antagonists (A-1155463 and A-1331852), we tested these agents against latency model cells generated from an additional 4 HIV-negative and 4 HIV-positive donors (**Table 1**). Treatment with A-1155463 alone drove statistically significant reductions in HIV DNA (1.88-fold average reduction, p=0.01) and IUPM (2.29-fold average reduction, p=0.05) (**Fig. 3F-G)**, as well as when combined with bryostatin-1 (2.24-fold average reduction in HIV DNA, p=0.02; and a 1.78-fold average reduction in IUPM, p=0.02) (**Fig. 3F-G).**Treatment with A-1331852 alone drove significant reductions in HIV DNA (2.15-fold average reduction, p=0.05) and IUPM (2.00-fold average reduction, p=0.03), and similar decreases when combined with bryostatin-1 (2.80-fold average reduction in HIV DNA, p=0.02; and a 1.99-fold average reduction in IUPM, p=0.02) (**Fig. 3H-I)**. Across these different experiments, the two BCL-X_L_ antagonists showed little in the way of cell toxicity, in contrast to the appreciable toxicity observed with the BCL-2 antagonist ABT-199 (**SFig. 1** & **Fig. 3**). Together, these results demonstrate that both of the BCL-X_L_ antagonists tested were sufficient to drive statistically significant reductions in a primary cell model of HIV latency, both with and without latency reversal by bryostatin-1.

**Table 1.**
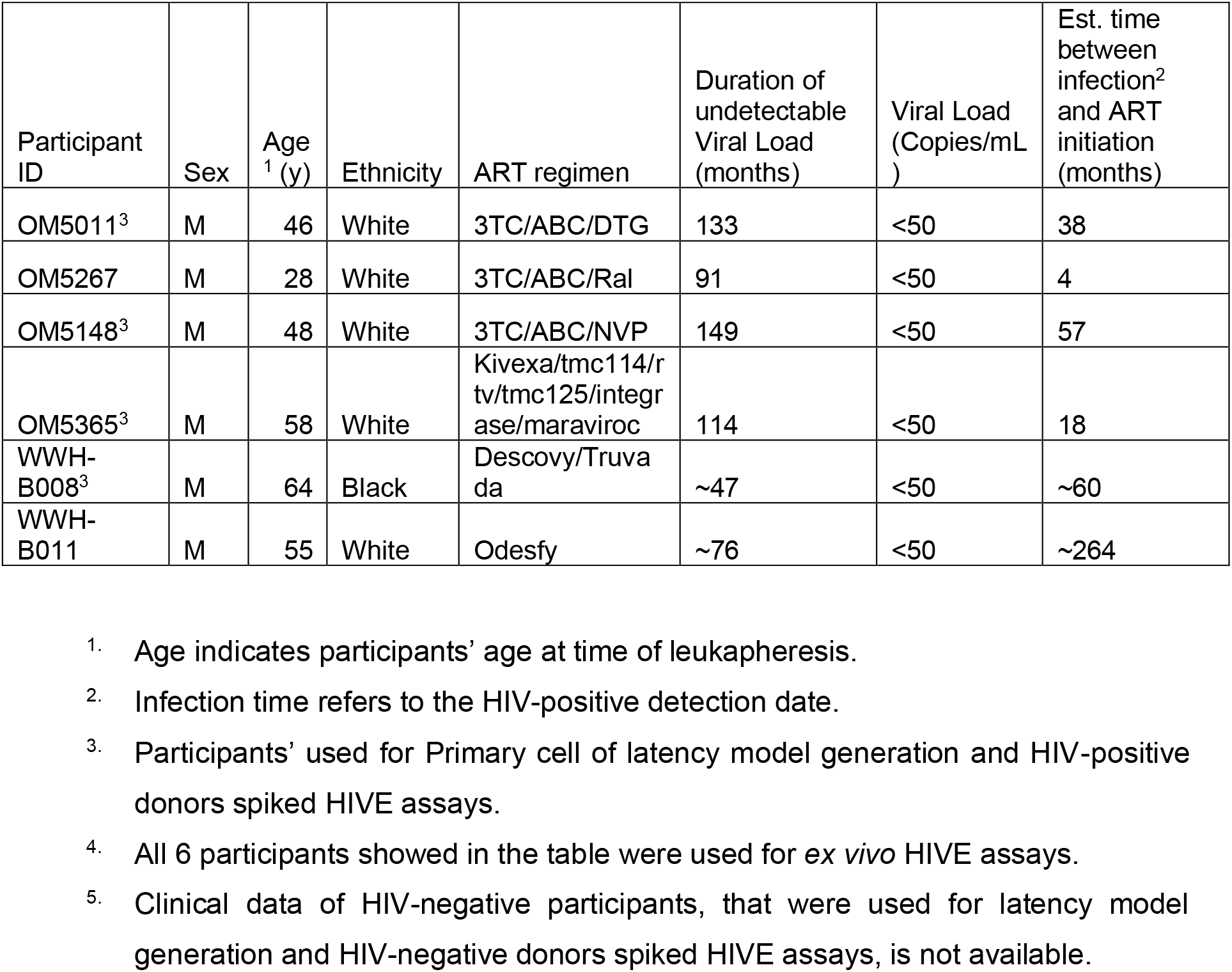
ART-suppressed participant clinical data.

### Selective BCL-X_L_ antagonists fail to reduce either total HIV DNA or infectious reservoirs from *ex vivo* ART-suppressed HIV-infected participants’ CD4^+^ T-cells

We next tested whether the reductions in infected cell frequencies observed with the BCL-X_L_ antagonist treatments in our primary cell latency model would also be recapitulated in HIVE assays targeting ‘natural’ HIV reservoirs in *ex vivo* CD4^+^ T-cells from ART-suppressed donors. A representative example of a HIVE assay performed with *ex vivo* samples is shown in **Fig. 4A-B**. In contrast to results from the primary cell model of latency, we observed a general lack of reductions in either HIV DNA or IUPM (**Fig. 4B**) for either of the BCL-X_L_ antagonists, used alone or in combination with bryostatin-1. The only significant differences that we observed were increases in IUPM following treatment with bryostatin-1 (p<0.001, **Fig. 4B**), which we have also reported previously [8]. Thus, this initial experiment with *ex vivo* CD4^+^ T-cells from a single donor showed a general lack of reduction in HIV-infected cell frequencies following treatment with BCL-X_L_ antagonists.

**Fig. 4.**
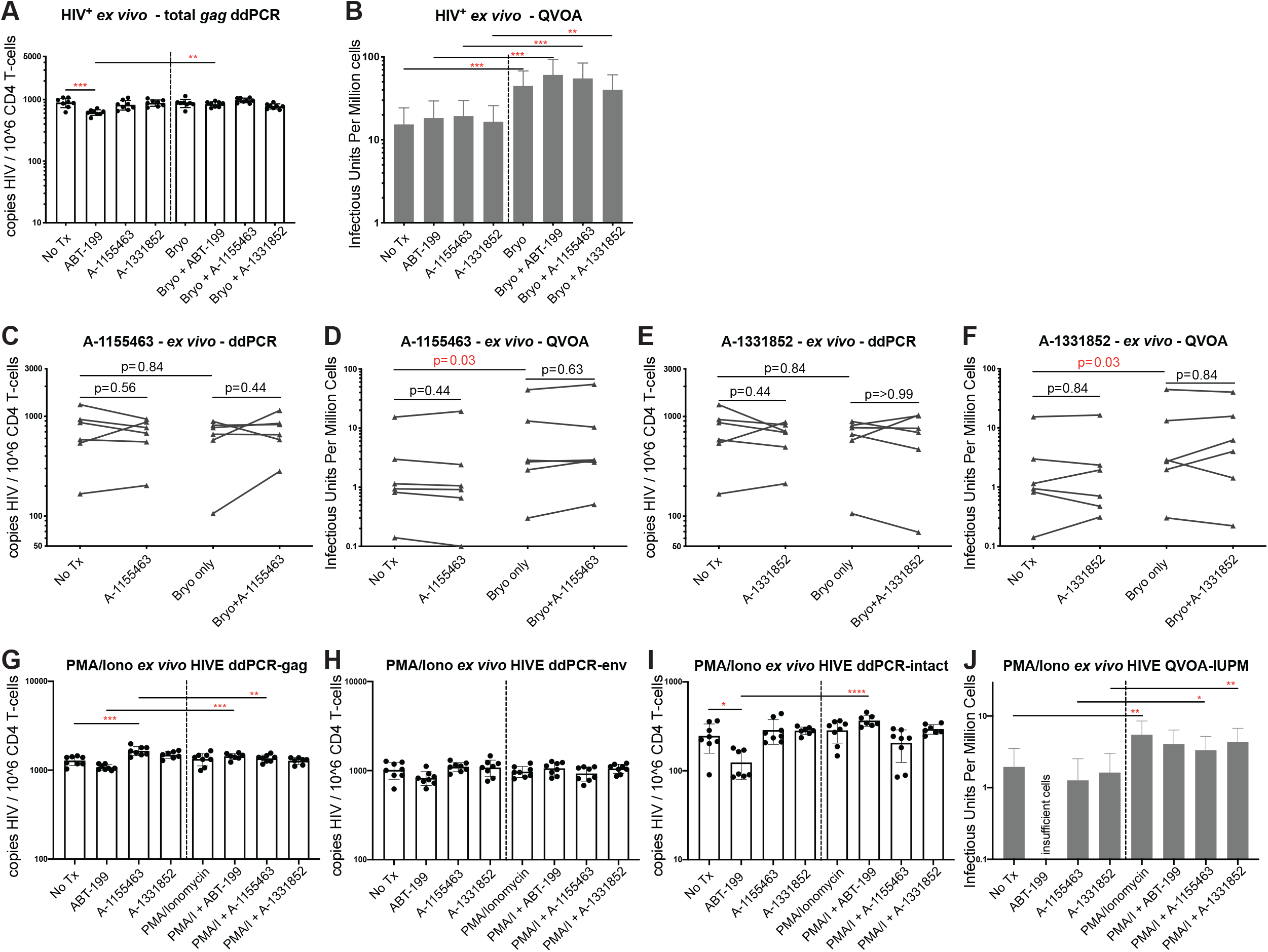
BCL-X_L_ antagonists failed to drive reductions in *ex vivo*, latently infected CD4^+^ T-cells in HIVE assays. HIVE assays were performed using *ex vivo* CD4^+^ T-cells from <u>ART-suppressed individual</u> WWH-B008. **(A)** ddPCR results using a 5’ end primer/probe set (HIV *gag*), for a representative HIVE assay. Shown are means ± SD of 8 replicates. **(B)** QVOA results from the same representative HIVE assay (WWH-B008), showing IUPM ± 95% confidence interval. Statistical significance determined by: one-way ANOVA for ddPCR, and a pairwise Chi-square test for QVOA (* p<0.05, ** p<0.01, *** p<0.001, **** p<0.0001). **(C)** Levels of HIV DNA (HIV *gag*) and **(D)** IUPM, comparing A-1155463 (100nM) vs No Tx, and Bryostatin-1+A-463 vs Bryostatin-1 (n=6). **(F)** Levels of HIV DNA (HIV *gag*) and **(F)** IUPM, comparing treatment with A-852 (100nM) vs No Tx, and Bryostatin-1+A-852 vs Bryostatin-1 (n=6). DMSO was added to NoTx conditions at a matched concentration with +Tx conditions. Statistical significance was determined by Wilcoxon matched-pairs signed rank test. HIVE assays were performed using *ex vivo* CD4^+^ T-cells from <u>ART-suppressed individual</u> WWH-B011. ddPCR results using a **(G)** 5’ end primer/ probe set (HIV *gag*), **(H)** 3’ end primer/probe set (HIV *env*), or **(I)** calculating double positives for a representative HIVE assay. Shown are means ± SD of 8 replicates. **(J)** QVOA results from the same representative HIVE assay (WWH-B011), showing IUPM ± 95% confidence interval. Statistical significance determined by: one-way ANOVA for ddPCR, and a pairwise Chi-square test for QVOA (* p<0.05, ** p<0.01, *** p<0.001, **** p<0.0001).

We extended these results by performing HIVE assays with *ex vivo* resting CD4^+^ T-cells from 6 long-term ART-suppressed HIV-positive donors (**Table 1**). Across this study population, we observed a lack of significant differences in either HIV DNA or IUPM when comparing untreated conditions to treatment with any of the BCL-X_L_ antagonists tested either alone or in combination with bryostatin-1 **(Fig. 4C-F)**, while the increase in IUPM observed with bryostatin-1 treatment in **Fig. 4B** was found to be consistent across this population (**Fig. 4D** & **F**). Additionally, we also tested the same concept in combination with a more potent LRA, PMA and ionomycin, and consistently observed the same lack of effects for the BCL-X_L_ antagonists. Again, we were not able to observe a significant difference in either HIV DNA, which was measured by *gag* primers/probe **(Fig. 4G**) or *env* primers/probe (**Fig. 4H)**, or “intact” provirus copies by calculating the double positives **(Fig. 4I)**, or IUPM **(Fig. 4J)** when comparing untreated conditions to treatment with any of the BCL-X_L_ antagonists tested either alone or in combination with PMA/ionomycin. Thus, in contrast to the primary cell latency model, neither of the BCL-X_L_ antagonists (A-1155463 and A-1331852) were sufficient to drive reductions in *ex vivo* viral reservoirs – including when combined with potent LRAs.

## Discussion

The intrinsic apoptotic pathway is one of the main contributors to CD4^+^ T-cell depletion during productive HIV infection [36–38]. BCL-2 and some of its family member proteins, including BCL-X_L_, can protect HIV-infected cells from apoptosis, and can contribute to the maintenance of the HIV reservoir under ART. Previously, we investigated the role of BCL-2 in helping to maintain HIV reservoirs under ART by conferring a survival advantage [8]. It is possible that BCL-2 family members are also involved in the establishment of HIV reservoirs. In the current study, we first tested the expression of pro-apoptotic and pro-survival BCL-2 family proteins during productive HIV infection by using an RNA-seq approach. We consistently observed the overexpression of the pro-apoptotic proteins Bak (*BAK1* coding protein), Bmf (*BM*F coding protein) and Noxa (*PMAIP1* coding protein), and the pro-survival proteins BCL-X_L_ (*BCL2L1* coding protein) and A1 (*BCL2A1* coding protein) in HIV-infected CD4^+^ T-cells, as compared to HIV-uninfected cells, for both total CD4^+^ and naïve-derived central memory CD4^+^ T-cells. We selected BCL-X_L_ for further investigation in the context of both productive and latent HIV infection based on these initial data and the availability of drugs in clinical development to target this molecule. BCL-X_L_ inhibits cell apoptosis by competing with the pro-apoptotic protein BAX/Bak [39–41]. In brief, as previously demonstrated by Kale et al., BID is first activated by caspase-8 mediated cleavage into cleaved BID (cBID), which is comprised of two fragments: BID-P7 and BID-P15; then, rapid high-affinity binding to cell membranes dissociates the P7 fragment into solution and favors insertion of the P15 fragment (tBID: truncated BID) into the membrane. Next, membrane-bound tBID recruits inactive BAX from the cytosol and then activates BAX, which inserts into the lipid bilayer, oligomerizing and permeabilizing the mitochondrial outer membrane thereby releasing intermembrane space proteins including cytochrome c and SMAC to induce cell apoptosis. When BCL-X_L_ is overexpressed, active tBID and BAX can recruit BCL-X_L_ to the membrane, resulting in inhibition of both pro- and anti-apoptotic proteins by mutual sequestration. BCL-X_L_ prevents tBID from activating BAX and prevents BAX from oligomerizing, resulting in the inhibition of mitochondrial outer membrane permeablisation (MOMP) [42]. The affinity of BCL-X_L_ is higher for tBID than for active BAX, and BAX-mediated membrane permeabilization is completely inhibited by BCL-X_L_ in sufficient quantities [43]. On the side of BAK, it was shown that BCL-X_L_ has a higher association affinity for BAK than for BAX [40], and BCL-X_L_ neutralizes BAK-mediated cell death - BAK is held in check only by Mcl-1 and BCL-X_L_, and can only induce apoptosis if freed from both [44]. In our RNA-seq results, both BCL-X_L_ and BAK were significantly overexpressed in the HIV-infected cells; thus, as BCL-X_L_ can prevent BAX-mediated apoptosis, this may lead to long-term survival of some of the HIV-infected cells, indicating a potential mechanism of HIV persistence.

The role of BCL-X_L_ during HIV-infection has not been well studied, but lessons can be learned from the setting of cancer, especially the study of solid tumors - where BCL-X_L_ overexpression is correlated with drug-resistance or cancer persistence. As BCL-X_L_ is essential for platelet survival [45–48], development and utilization of BCL-X_L_ antagonists need to be evaluated for safety and off-target toxicity. In high-risk B-cell acute lymphoblastic leukemia (B-ALL), loss of functional IKAROS and increased expression of CK2 results in increased expression of BCL-X_L_, which is associated with resistance to doxorubicin treatment [49]. In the differentiated thyroid carcinomas (DTC) diagnosis, BCL-X_L_ expression was found to be high, and was confirmed as an independent prognostic factor for persistent disease [50]. BCL-X_L_ is also a key survival factor in activation of stromal fibroblasts (ASFs) as well as in senescent cholangiocytes. Treatment with the BCL-X_L_-specific inhibitor A-1331852 reduces liver fibrosis, possibly by a dual effect on activated fibroblasts and senescent cholangiocytes [51]. In the condition of ER-stress, STING induction was enhanced in EGFR drug tolerant persister cells, and this also link to a BCL-X_L_ dependent mitochondrial protection and cell survival in the absence of ufmylation. Cell death following ER-stress induction in EGFR TKI persister cells could be triggered by using the BCL-X_L_ inhibitor A-1331852 [52]. *In vivo*, selective inhibition of BCL-X_L_ (by WEHI-539 or A-1155463), but not BCL-2, resulted in a decrease in tumor growth rate in an orthotopic Swarm Rat Chondrosarcoma (SRC) model [53]. A-1155463 was tested *in vivo* and showed a reversible thrombocytopenia in mice and inhibited H146 small cell lung cancer xenograft tumor growth following multiple doses [54]. A-1155463 and A-1331852 showed the potential to enhance the efficacy of docetaxel in solid tumors and avoid the exacerbation of neutropenia [55]. In addition to this, the BCL-X_L_ antagonists were also tested as senolytic agents, and induced apoptosis in senescent cells [56]. Given the overexpression of BCL-X_L_ in HIV infected cells in our study, and the character of HIV latency, this evidence suggests that the BCL-X_L_ may also contribute to HIV-infected cell persistence and may even drive infected cells into latency, which warrants further investigation.

Given the potential of BCL-X_L_ to contribute to HIV persistence, we were interested in testing whether BCL-X_L_ antagonists are able to eradicate productively HIV-infected cells, as well as long-lived HIV reservoir-harboring cells. In *in vitro* productive infection, both of the selective BCL-X_L_ antagonists tested showed some efficacy in infected cell killing, which was specific for HIV-infected cells (where BCL-X_L_ expression was upregulated). Thus, we moved to a primary cell model of HIV latency [33]. Selective BCL-X_L_ antagonists were used alone (sensitize to cell apoptosis without cell activation) or in combination with a latency reversing agent to reactivate virus and promote viral cytopathic effects (kick and kill). Since this primary cell model of HIV latency was mainly used to test the concept of the antagonists against the target cells only – whether they can sensitize the HIV reservoir-harboring cells to apoptosis, no immune-effector cells were added. We observed significant reductions in HIV infected cell frequencies, both total HIV DNA level and replicative competent virus level (IUPM), providing initial support for the concept of utilizing BCL-X_L_ antagonists for HIV cure strategies.

This prompted us to test this strategies against HIV reservoirs in *ex vivo* CD4^+^ T-cells, representing long-lived cells potentially selected by immune and other pressures. Similar to our recent work with BCL-2 antagonism ABT-199 [8], the reductions in infected cells observed in the latency model system did not extend to measurable reductions in *ex vivo* reservoir, whether BCL-X_L_ antagonists were used alone or in combination with Bryostatin-1. While the reasons underlying this difference are currently unclear, they fit a pattern in our recent results, which have indicated that success in various kick and kill combinations that are effective against primary cell models of latency, have not translated into success against reservoirs in *ex vivo* CD4^+^ T-cells from ART-treated donors: including CTL + LRAs [57], BCL-2 antagonists + LRAs [8], and now BCL-XL antagonists +/− LRAs. While additional research in this area is needed, the results fit a model that we have recently proposed which postulates that long-lived reservoir harboring cells in people living with HIV have been selected not only with respect to proviral latency, but also with respect to the intrinsic survivability of the reservoir harboring itself (reviewed in [58]). Either or both of these factors may have contributed to our results, where we acknowledge that limitations in latency reversal in *ex vivo* reservoirs may have also limited the impact of BCL-X_L_ antagonists, even in the case of PMA/I stimulation. Our results call for further study into these mechanisms. We suggest that BCL-X_L_ antagonists will be a useful tool in this regard given their: ease of use, low toxicity to bystander cells, and clear dichotomy between the model and *ex vivo* system.

Finally, we would propose the overall interpretation of our results as a partial success of BCL-X_L_ antagonists, encouraging of their further testing and development. The ability to drive reductions in *ex vivo* HIV reservoirs appears to be a ‘high bar’ to clear, and we interpret activity against the latency model as motivation for future studies testing BCL-X_L_ antagonists in combination with: immune effectors such as CTL (shown to be partially effective in combination with BCL-2 antagonists), novel LRA combinations, and other emerging agents which show promise in terms of overcoming remaining barriers to the elimination of reservoir-harboring cells.

## Materials and Methods

### Agents: Latency reversing agents, Chemical agents and Antibodies

LRAs and BCL-X_L_ antagonists were used at the following concentrations: Bryostatin-1 dissolved in DMSO and used at 10nM (Sigma-Aldrich); PMA and Ionomycin (both Sigma-Aldrich) dissolved in DMSO and used at 50ng/mL and 1μM, respectively; anti-CD3 (OKT3, Biolegend) and anti-CD28 (CD28.2, Biolegend) antibodies used at 1μg/mL each; ABT-199 (Med Chem Express, Cat# HY-15531), A-1155463 (Med Chem Express, Cat# HY-19725), and A-1331852 (Med Chem Express, Cat# HY-19741) were dissolved in DMSO and used at 100nM each. Antibodies for cell staining included: fixable viability dye (aqua, ThermoFisher), anti-human CD3 (clone SK7, Biolegend, 1:200), anti-human CD4 (clone RPA-T4, BD Biosciences, 1:200), anti-human CD8 (clone RPA-T8, Biolegend, 1:200), anti-human CD45RA (clone HI100, Biolegend, 1:200), anti-human CCR7 (clone G043H7, Biolegend, 1:200), anti-human CD69 (clone FN50, Biolegend, 1:200), anti-human HLA-DR (clone L243, Biolegend, 1:200), anti-human BCL-X_L_ (clone 7B2.5, Invitrogen, 1:50), and p24 antibodies (anti-HIV core antigen: clone KC57, Beckman Coulter, 1:200).

### HIV infection and flow cytometry staining for BCL-X_L_ expression

Total CD4^+^ T-cells were enriched from HIV-negative donor PBMCs by magnetic negative selection, following the manufacturer’s instructions (StemCell Technologies). These cells were then activated with 1μg/mL each of anti-CD3 and anti-CD28 antibodies in RPMI-10 media supplemented with 50IU/mL of IL-2 (R10-50). Activated CD4^+^ T cells were infected with HIV strain (‘JR-CSF’, ‘NL4-3’, or supernatant collected from a QVOA p24-positive well from donor OM5334 [named as OM5334 QVOA virus]) by spinoculation at 2000xg for 2 hours. Cells were then incubated in R10-50 at 37°C for 7 days, with half medium changes every other day. Cells were collected and treated with DMSO (1:1000 as control), ABT-199, A-1155463 or A-1331852 in R10-50 for 48 hours. Cells were then stained with antibodies against human-CD3, CD4, CD8, BCL-X_L_ and p24, with a fixable live/dead dye staining, and then analyzed by flow cytometry (Attune NxT). Data were analyzed using FlowJo software.

### RNA-seq sample acquisition

Cultured T_CM_ CD4^+^ T-cells were generated as previously described [32], also see below. Total CD4^+^ T-cells or T_CM_ CD4^+^ cells were activated and infected as described above. When the infection rate went over 10%, cells were collected and stained with antibodies against human- CD3, CD4, CD8, and p24, with a fixable live/dead dye staining, and then sorted by flow cytometry (SONY9000) directly into 15mL collection tubes for HIV+ (p24^+^) and HIV-(p24^-^) cells (**Fig. 1A**). Total RNA was immediately extracted using the miRNeasy FFPE Kit (Qiagen), and RNA quality and concentration was determined by Agilent Bioanalyzer 2100. Library preparation was performed using the methods of TruSeq RNA Sample Preparation (Non-Stranded and Poly-A selection), and sequencing was run on HiSeq4000 (Illumina) with a single read clustering and 100 cycles of sequencing.

### RNA-seq data analysis

The raw sequencing reads in BCL format were processed through bcl2fastq 2.19 (Illumina) for FASTQ conversion and demultiplexing. Reads were aligned with default parameters to the human reference genome (GRCh38.p12) with STAR (ver. 2.6.0c) [59]. Gene abundance was calculated with featureCounts using composite gene models from Gencode release 28. Differentially expressed genes were determined with DESeq2 using Wald tests (q < 0.01) [60]. Expression heatmaps of the genes encoding proteins of the BCL-2 family were generated using variance-stabilized data, with the values being centered and scaled across each gene. The code and read count matrix are available upon request.

### Generation of “cultured T_CM_” CD4+ primary T-cells

Latency model cells were generated as previously described [32]. 5×10^6^ naïve CD4^+^ T cells from HIV^−^ or HIV^+^ donors were isolated by magnetic negative selection (StemCell Technologies), then cultured at 10^6^ cells/mL in 96-well plates using R-10 media (RPMI-1640 media supplemented with 10% FBS, 2mM L-glutamine, 100 units/ml Penicillin and 100μg/ml Streptomycin) supplemented with 12.5uL/mL of dynabeads human T-activator CD3/CD28 (Invitrogen), 2μg/mL anti-human IL-12 (PeproTech), 1μg/mL anti-human IL-4 (PeproTech), and 10ng/mL of TGF-β1. After 3 days, dynabeads were removed by magnetic selection and cells were washed, followed by culture in R-10 media supplemented with 30 IU/mL of IL-2 (R-10-30). Media was changed on days 4 and 5 with fresh R-10-30 media.

### Infection of “cultured T_CM_” cells to generate latency model

On day 7 of the above “cultured T_CM_” generation protocol, 1/5th of the cells were infected with HIV_NL4-3_ at a MOI of 0.6 by spinoculating at 2000 x g for 2 hours, at 37°C. Cells were then resuspended in R-10-30 with the other 3/5th of the cells and placed back in the incubator, while the remaining 1/5th were set aside as an uninfected control. On day 10, cells were recounted, washed and resuspended in R-10-30 at 1M cells/mL and plated in 96-well round bottom plates to crowd infection. On day 13, cells were washed and resuspended at 1M cells/mL, and then transferred to culture flasks in R-10-30 supplemented with 1μM Raltegravir and 0.5μM Nelfinavir at 1M cells/mL. On day 17, CD4 positive cells were isolated by magnetic positive selection following the manufacturer’s instructions (Life Technologies), and then used in the various assays. A small portion of the cells were reactivated with CD3/CD28 dynabeads and stained for intracellular Gag, to determine infection percentages and ensure quality controls.

### “cultured T_CM_” primary cell latency model “spiked” HIV Eradication (HIVE) Assays

HIVE assays were set up as previously described [57]. Briefly, >15M CD4^+^ T cells were pulsed with or without bryostatin-1 for 2 hours, then washed and co-cultured with or without either BCL-X_L_ antagonists (A-1155463 or A-1331852) in XVIVO-15 media (Lonza) supplemented with 1μM Tenofovir Disoproxil Fumarate, 1μM nevirapine, 1μM emtricitabine, 10μM T-20, 10U/ml human DNAse I (ProSpec), and 0.1nM IL-7 (HIVE media). Primary latency model cells were spiked into newly isolated, autologous resting CD4^+^ T-cells to achieve a frequency of ~1,000 – 10,000 copies of HIV DNA per million CD4+ T-cells. Following a 4 days co-culture, CD4^+^ T cells were isolated and rested for 24 hours in R10-50 media at 37°C to allow for an ARV washout period. Aliquots of pre- and post-CD4 enrichment samples were collected and stained for viability and memory phenotype/activation status with antibodies against CD3, CD4, CD8, CD45RA, CCR7, CD69 and a viability dye (Invitrogen Technologies), then analyzed by flow cytometry. Following the overnight culture, a small aliquot of cells was mixed with CountBrightTM absolute counting beads and viability dye to obtain a count of total, live CD4 ^+^ T-cells by flow cytometry. This viable cell count was used to determine cell numbers for ddPCR and QVOA plating strategies (**Fig. 3A**).

### *Ex vivo* HIV reservoir Eradication (HIVE) Assays

HIVE assays with *ex vivo* resting CD4^+^ T cells from long-term ART- suppressed donors were set up same as described above, without “cultured TCM” primary cell latency model “spiked” in.

### Digital droplet PCR

ddPCR measuring total HIV DNA (HIVEs) was performed as previously described [61], with slight modifications. For each PCR reaction, 5 units of restriction enzyme BsaJI (NEB) was directly mixed with 300ng of DNA, ddPCR Supermix (no dUTP) for Probes (Bio-Rad), and final concentrations of 900nM primers and 250nM probe. Primers/Probes were listed in **Table 2**; Droplets were prepared using the QX200 Droplet Generator (Bio-Rad) following the manufacturer’s instructions. Sealed plates were cycled using the following program: 95°C for 10 min; 40 cycles of 94°C for 30 s, 60°C for 1 min; and 98°C for 10 min. Reactions were analyzed using the QX200 Droplet Reader and number of template molecule per μl of starting material was estimated using the Quantalife ddPCR software. 8 technical replicates were run per sample, and we consistently applied a pre-determined exclusion criterion to outliers that deviated from mean values by >2x the standard deviation.

**Table 2.**
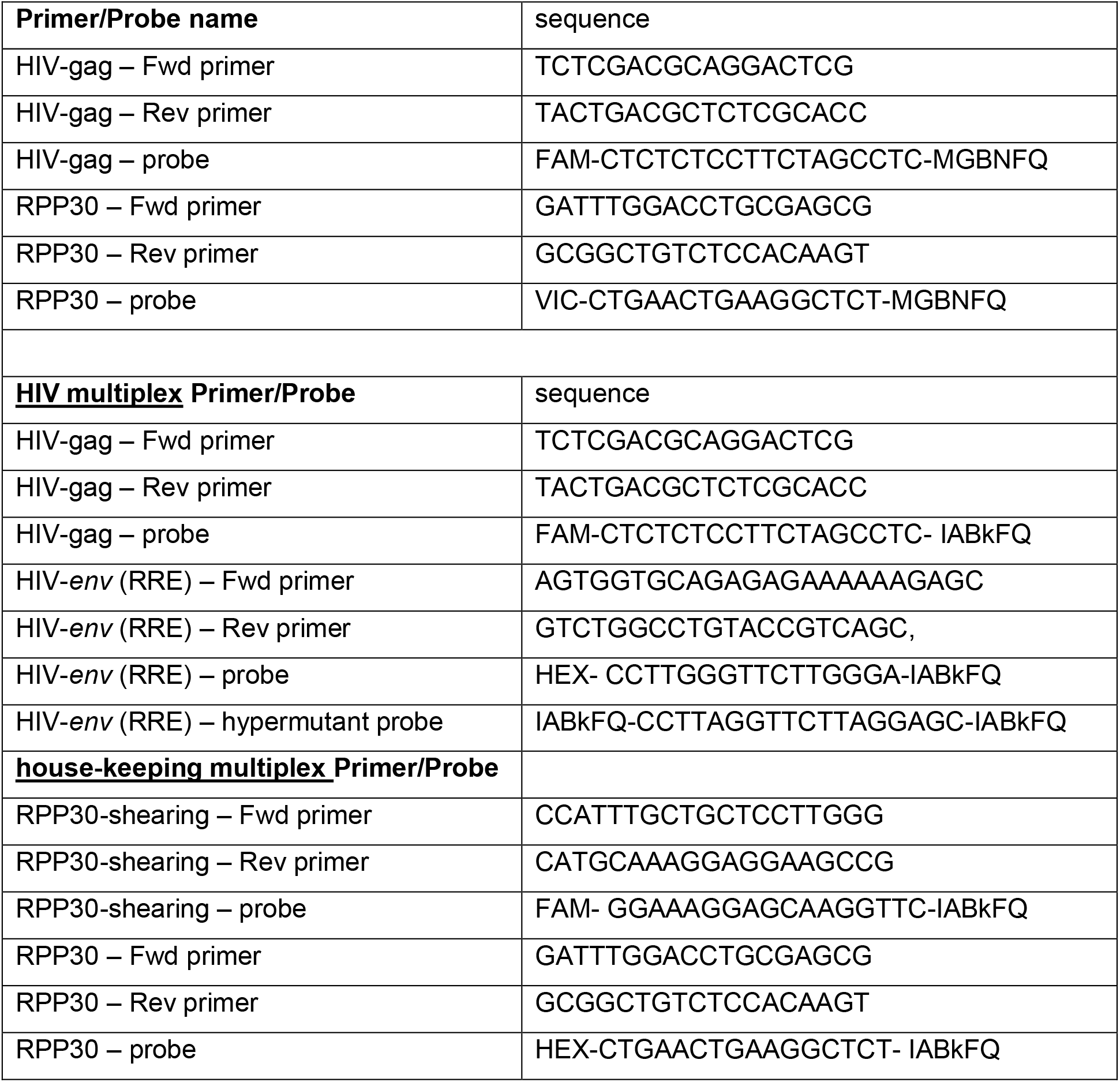
Primers and probes used for ddPCR.

For HIVE in Fig. 4G-I, a modified IPDA [62] was applied. For each PCR reaction, same ddPCR supermix and final concentrations of primers and probes as above, but with 5 units of restriction enzyme Xho I (NEB) mixed with 750ng DNA. Primers and probes were used in 2 separate PCR systems: 1) <u>house-keeping multiplex</u> with RPP30 and RPP30-shearing; 2) <u>HIV multiplex</u> with *gag* and HIV-*env* (RRE). Primers/Probes were listed in **Table 2**. PCR program is as following: 95°C for 10 min; 45 cycles of 94°C for 30 s, 53°C for 1 min; and 98°C for 10 min. DNA input of house-keeping multiplex, is 100-fold diluted from the input of HIV-multiplex. Total *gag, env* or ‘intact’ proviruses copies were calculated by multiplying the dilution factors, and ‘Intact provirus’ copies were corrected with the shearing percentage calculated from house-keeping multiplex. Same as above, 8 technical replicates were run per sample, and applied with a pre-determined exclusion criterion to outliers that deviated from mean values by >2x the standard deviation.

### Quantitative viral outgrowth assays (QVOAs)

QVOAs were performed using a previously described protocol [63], with slight modifications depending on the application. Live cells counted by flow cytometry were distributed into either three of 2-fold serial dilutions with 12 replicates per dilution. This was determined based on the numbers of viable cells recovered at the end of each HIVE assay and the baseline IUPM values of the donor. Cells were then stimulated with 2μg/ml of PHA (ThermoFisher Scientific) + 1M PBMCs (HIV-negative donor, irradiated at 5000 rads). The next day, 1M CCR5^+^MOLT-4 cells were added along with a half media change. Cultures were then incubated for 14 days, with half media changes with R10-50 every 3-4 days. We performed p24 ELISA on supernatant 15d after the PHA stimulation. For each condition, values for cells/well, number of positive wells, and total wells tested were entered into a limiting dilution analyzer (http://bioinf.wehi.edu.au/software/elda/) [64] to calculate the maximal likelihood IUPM and a corresponding 95% confidence interval.

### Quantification and Statistical Analysis

Statistical analyses were performed using Prism 8 (GraphPad), and the statistical analysis methods used are reported in the Figure Legends. All ddPCR data were analyzed by ordinary one-way ANOVA, with Tukey’s multiple comparisons test if the overall ANOVA F-test was significant; statistics for the summary datasets for HIV DNA were performed using the mean of 8 replicates per condition. QVOAs were run at the end of each HIVE assay, and the IUPM was calculated as described above; pairwise comparisons were made by a Chi-square test by the method of Hu and Smyth (2009) [64] using ELDA software (http://bioinf.wehi.edu.au/software/elda/). All comparisons between HIVE conditions used paired non-parametric test (2-tailed) - Wilcoxon matched-pairs signed rank test. A p-value of less than 0.05 was considered significant.

### Study Approval

People living with HIV were recruited from either the Maple Leaf Medical Clinic in Toronto, Canada through a protocol approved by the University of Toronto Institutional Review Board (IRB), or Whitman-Walker Health in Washington D.C. (**Table 1**). Additional use of de-identified samples was reviewed and approved by the George Washington University (Washington, D.C.), and Weill Cornell Medicine (New York) Institutional Review Boards. All subjects were adults, and gave written informed consent prior to their participation. Leukapheresis samples were used immediately if possible, or cryopreserved in liquid nitrogen; cells were not left in culture prior to the initiation of experiments.

## Acknowledgments

This work was supported by the NIH funded R01 awards AI31798, AI147845, and the R56 award AI52764. It was also supported in part by the Martin Delaney ‘BELIEVE’ Collaboratory (NIH grant 1UM1AI26617); and the NIH funded Center for AIDS Research grants (P30 AI117970), which are both supported by the following NIH Co-Funding and Participating Institutes and Centers: NIAID, NCI, NICHD, NHLBI, NIDA, NIMH, NIA, FIC, and OAR. The following reagents were obtained from the NIH AIDS Research and Reference Reagent Program: IL-2, pNL4-3, CCR5^+^ MOLT-4 cells. Reagents for HIV p24 ELISAs were obtained from the NCI’s AIDS and Cancer Virus Program.

## Author Contributions

RBJ and YR conceptualized the study. RBJ, YR, SHH and AB developed the methodology. RBJ, YR, SHH, ABM, WDCA, TK, LL, DC, RT and TR conducted the investigation. PZ, FD and DB provided bioinformatic analysis for the RNA-seq data. CK, EB, AW, CC, and WDH recruited study participants and provided clinical samples. RBJ and YR wrote the original draft of the manuscript, with edits provided by all co-authors. RBJ acquired funding. RBJ and AB supervised the study.

## Declaration of Interests

YR, SHH, & RBJ declare that they are inventors on a patent that claims the use of BCL-2 and BCL-X_L_ antagonists as therapies for HIV. RBJ declares that he has received payments for his role on the scientific advisory board of AbbVie Inc. WDH declares payment for scientific advisory boards and consulting services for CSL Behring (Calimmune), Enochian Biosciences, Gilead, Merck, ViiV/GSK.

---

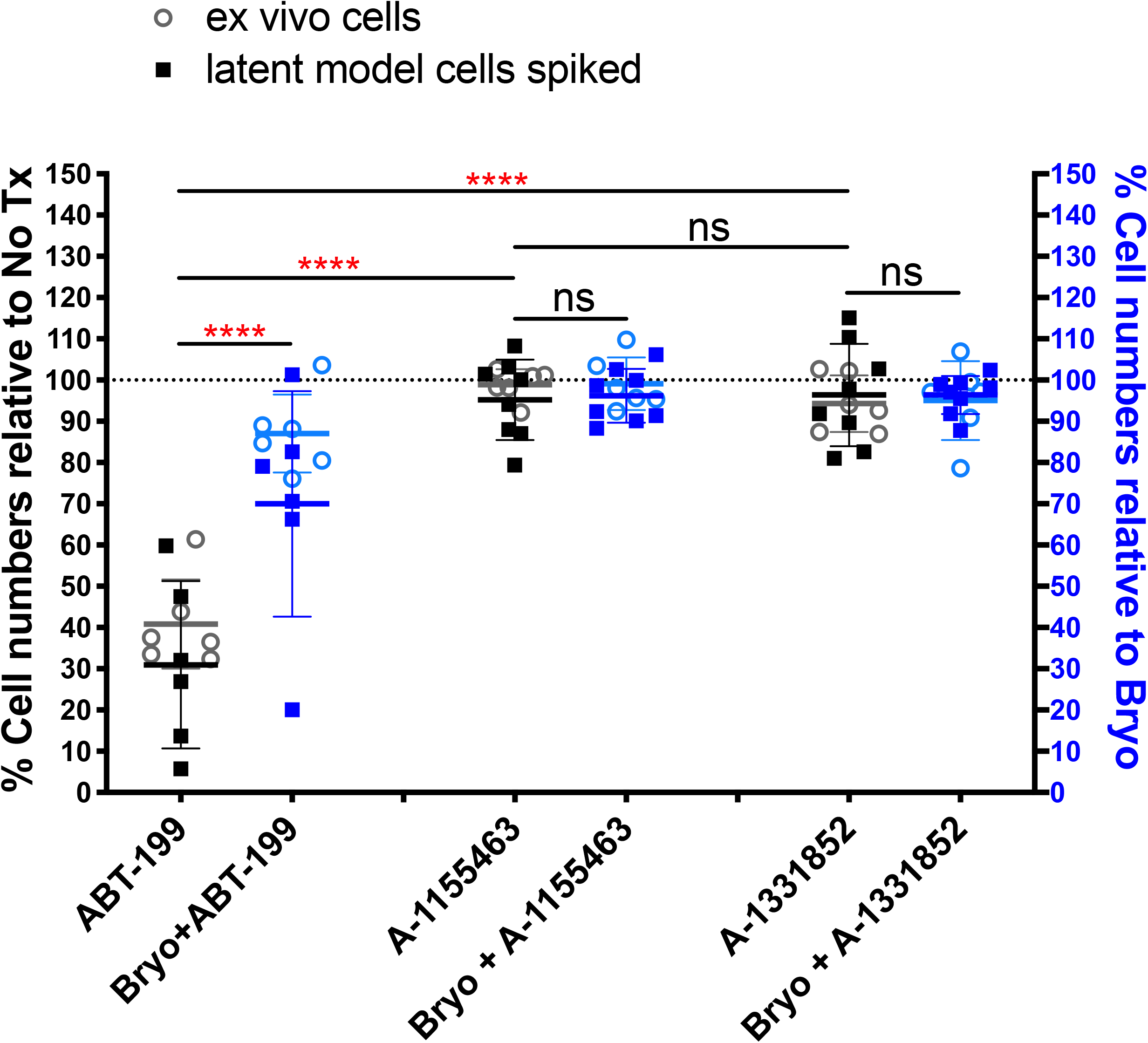

